## Supplementary material for "Network-based clustering unveils interconnected landscapes of genomic and clinical features across myeloid malignancies"

#### A Simulations

For our setting, we consider a set of variables representing the mutation profiles of patient tumours, which we refer to as mutations, and a set of clinical covariates for each patient, which we refer to simply as covariates. To evaluate the performance of our covariate-adjusted clustering method, we benchmarked it against standard clustering methods over a range of different simulations, which were selected to cover typical pan-cancer applications. To account for our biological assumptions, we simulated data with cluster-specific probabilistic relationships and causal effects from covariates on the clustered variables. We simulated a Bayesian network for each cluster, including covariates as nodes with outgoing edges and constant conditional probability tables.

We compared our method against the following clustering algorithms: a Bayesian network mixture model (BNMM), k-means and a Bernoulli Mixture Model (BMM). Since these clustering algorithms do not adjust for covariates, we applied them once by adding the covariates to the clustered variables and once by excluding the covariates from the analysis. Code implementing our method and reproducible simulation benchmarks are available at `https://github.com/cbg-ethz/myeloid-clustering`.

Figure S1 shows the benchmark results over different numbers of covariates, where clustering accuracy was assessed via the adjusted rand index (ARI). In the presence of covariates, the most accurate clustering performance is reached by our covariate-adjusted method. This is in line with our theoretical expectations, which removed the effect of the covariates on the cluster membership probability while adjusting for their other effects. In general, the network-based clustering methods have a higher clustering accuracy, which reflects that these structures allow one to model probabilistic relationships among the variables. Figure S1 shows that k-means and BMM perform better when the covariates are excluded from the analysis. This can be explained by the fact that these models assume independent probability distributions and hence including the covariates only increases the variance in the cluster assignment. In contrast, the performance of the BNMM improves when including the covariates, since their induced probabilistic relationships allow one to model the data more accurately.

To assess the performance of our covariate-adjusted algorithm in more detail over a range of different scenarios in our benchmark, we varied the following cluster-specific parameters: number of clusters $K$, number of variables $N_V$, number of covariates $N_C$, and number of samples per cluster $n_k$. While varying one parameter in our benchmark, all other parameters remained constant. The results of the different benchmarks are displayed in Figure S2.

#### B Application to TCGA Pan-Cancer Data

We applied our clustering algorithm to a large-scale genomic dataset from the Cancer Genome Atlas (TCGA)[1], which includes the mutational profiles and clinical information of 8085 patients from 22 different cancer types. For each primary cancer type, we considered the 16 most significantly mutated genes, adding up to a total of 201 genes across all cancer types[2]. In addition, we included the clinical covariates age, sex, and cancer type in the analysis. Since the variables age and sex have a downstream causal effect on the mutations, we adjusted for them in our covariate-adjusted clustering framework. The number of clusters was determined by calculating the Akaike information criterion (AIC) for a range of different sizes.

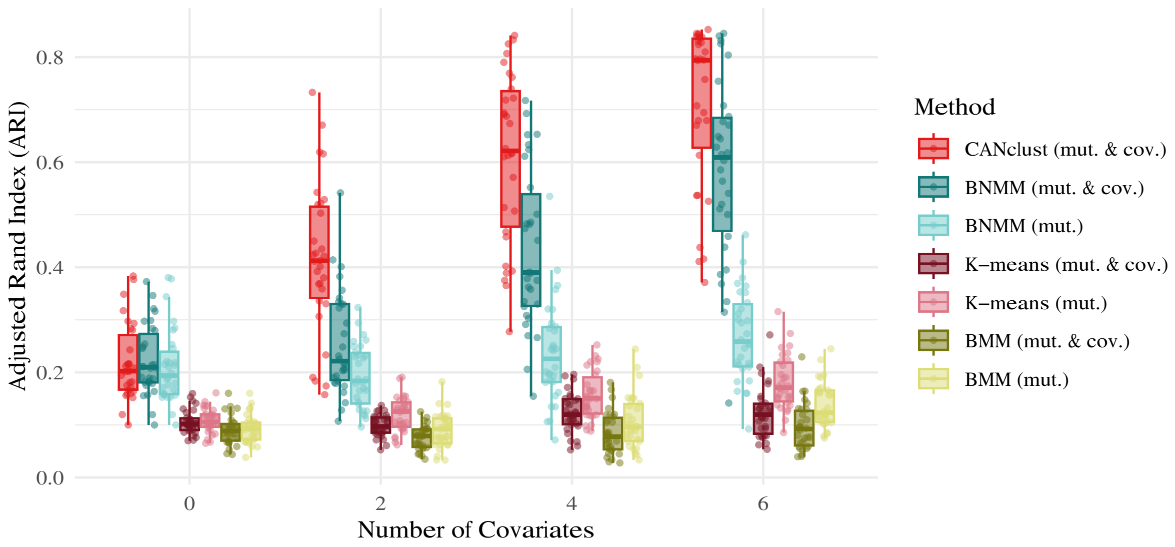


**Figure S1.** Simulation benchmark for an increasing number of covariates, comparing our covariate-adjusted network clustering algorithm (CANclust) against a Bayesian network mixture model (BNMM), k-means, and a Bernoulli mixture model (BMM). Standard clustering algorithms that do not adjust for covariates were applied twice: first, by including the covariates, and second, by excluding the covariates.

While there is no ground truth cluster assignment in pan-cancer data, similarity in survival outcomes within the clusters is a proxy indicator of shared biological properties. We applied the Cox proportional hazards regression model and corrected for the effects of age, stage, and cancer type on the results since these are strong predictors of survival.

Using our method to cluster the mutational profiles based on their probabilistic relationships, we identified 22 novel cancer subgroups for which the survival probabilities are shown in Figure S3. Each subgroup is associated with a distinct network, representing its subgroup-specific probabilistic relationships. Table S1 shows the corrected likelihood ratio (LR) of the Cox proportional hazards regression model for different clustering algorithms, where we only considered network-based approaches due to their superior performance in the benchmark study. High corrected LRs indicate cluster assignments that are highly predictive of survival. The lowest corrected LR is reached by the BNMM that neglects the clinical covariates, highlighting the importance of integrating these when clustering mutational profiles. In contrast, the cancer subgroups found with our covariate-adjusted clustering method were most predictive in survival beyond clinical information (LR $= 46.6$, p-value $= 1.0 \times 10^{-10}$), confirming our theoretical expectations and simulation results.

**Table S1.** Likelihood ratio for different clustering algorithms after accounting for clinical and histopathological covariates in the survival analysis.

| Method | Corrected likelihood ratio | P-value |
|---|---|---|
| CANclust | **46.6** | $1.0 \times 10^{-10}$ |
| BNMM (mut. & cov.) | 43.8 | $4.0 \times 10^{-10}$ |
| BNMM (mut.) | 34.4 | $5.6 \times 10^{-07}$ |

### C Supplementary Methods

#### C.1 Adjusting for Covariates

We want to cluster the mutational profiles to a number of $K$ individual clusters. We assume that the mutational profiles $X_V$ can be represented by the probability distributions of $K$ Bayesian networks, and that the cluster-dependent clinical covariates $X_C$ affect mutations via directed edges from covariates to mutations.

In the Bayesian network mixture model, we can consider the following three clustering variations: covariate-adjusted clustering, joint clustering of mutational profiles and clinical covariates, and clustering of mutational profiles only.

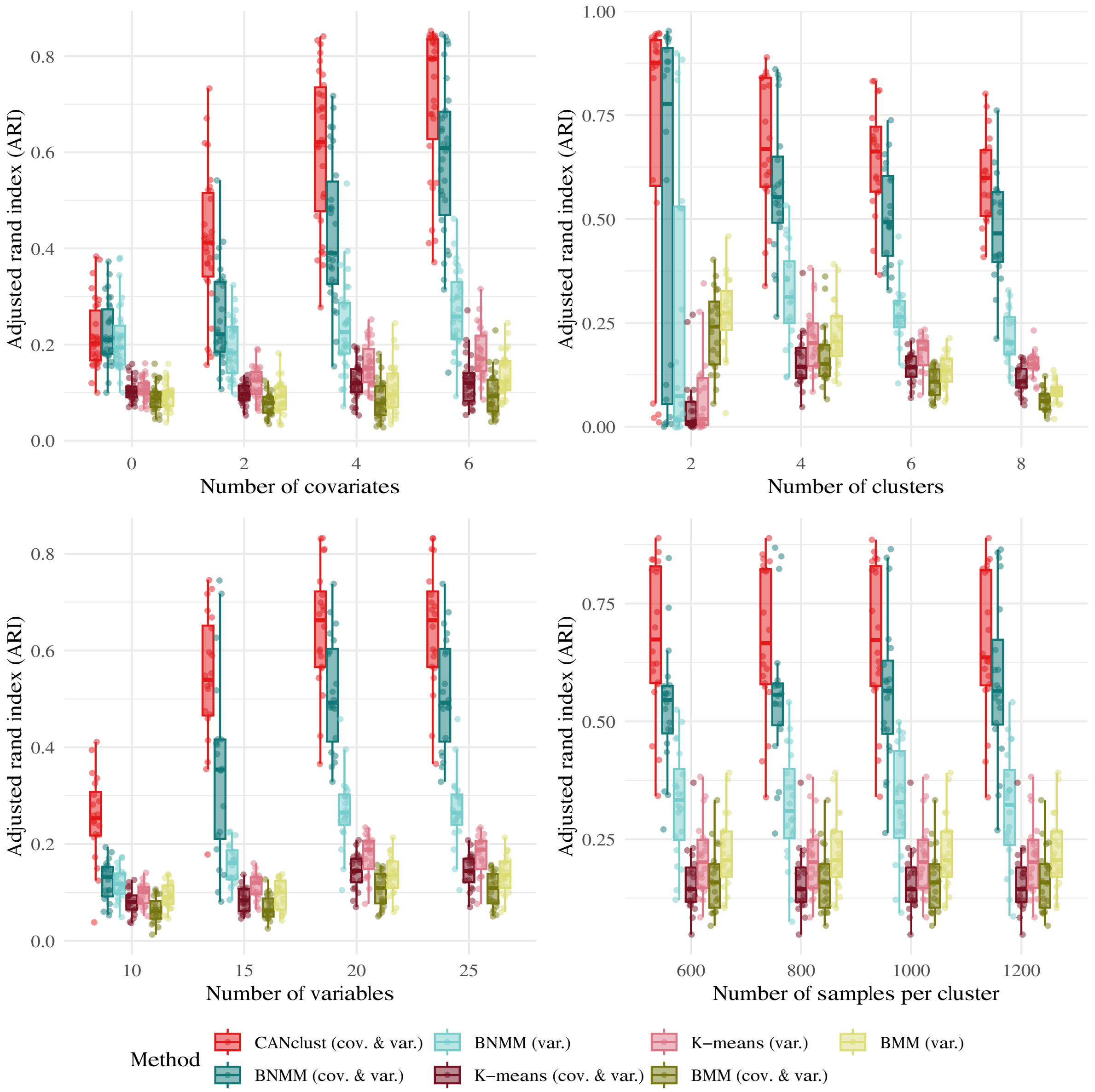


**Figure S2.** Simulation benchmark over a range of varied cluster-specific parameters including our covariate-adjusted network clustering algorithm (CANclust), a Bayesian network mixture model (BNMM), k-means, and a Bernoulli mixture model (BMM).

For the covariate-adjusted clustering, the membership probability function is defined as

$$\tilde{\phi}(X_V \mid k) = \frac{\gamma_k \cdot P\left(X_V \mid X_C, \hat{\mathscr{G}}_k, \hat{\theta}_k\right)}{\sum_{k'=1}^{N} \gamma_{k'} \cdot P\left(X_V \mid X_C, \hat{\mathscr{G}}_{k'}, \hat{\theta}_{k'}\right)} \,. \tag{1}$$

For the clustering of mutational profiles and clinical covariates, it is

$$\phi(X_V, X_C \mid k) = \frac{\gamma_k \cdot P\left(X_V, X_C \mid \hat{\mathscr{G}}_k, \hat{\theta}_k\right)}{\sum_{k'=1}^{N} \gamma_{k'} \cdot P\left(X_V, X_C \mid \hat{\mathscr{G}}_{k'}, \hat{\theta}_{k'}\right)} \,. \tag{2}$$

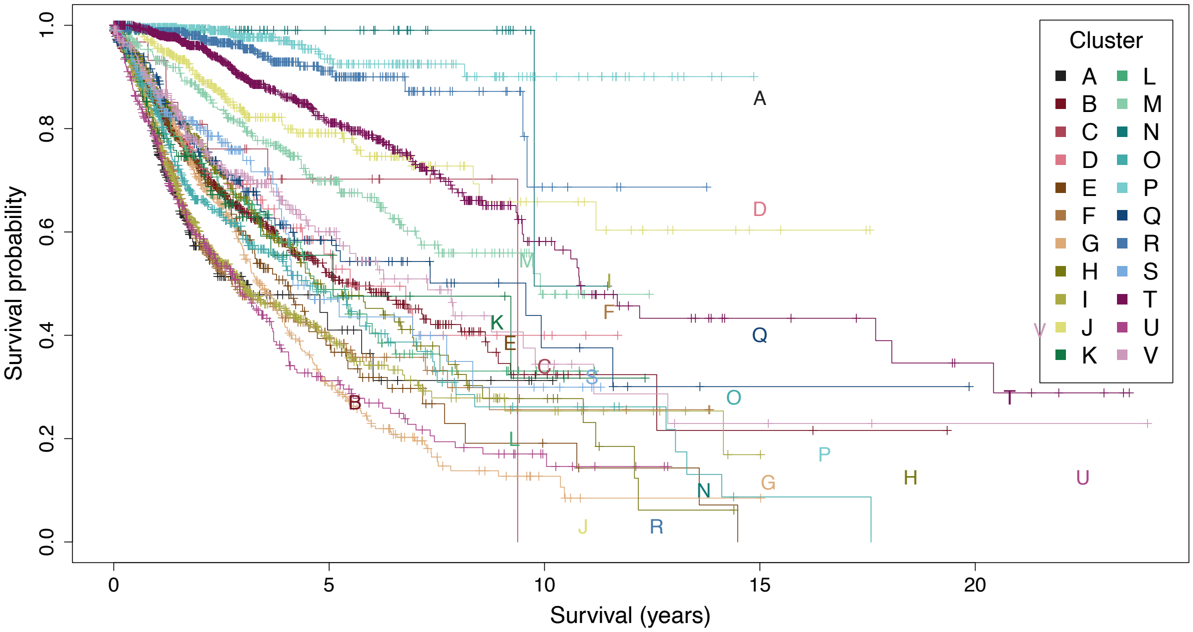


**Figure S3.** Survival probabilities of the novel cancer subgroups that were learned using network-based clustering, labelled A-V.

For the clustering of mutational profiles, it is

$$\phi(X_V \mid k) = \frac{\gamma_k \cdot P\left(X_V \mid \hat{\mathscr{G}}_k, \hat{\theta}_k\right)}{\sum_{k'=1}^{N} \gamma_{k'} \cdot P\left(X_V \mid \hat{\mathscr{G}}_{k'}, \hat{\theta}_{k'}\right)} \,. \tag{3}$$

If we assume that the probability distribution of the cluster-independent covariates $X_C$ is constant across different clusters, i.e. $\forall k, k' \in \{1, ..., K\} : P(X_C \mid k) = P(X_C \mid k')$ and if the Bayesian network parameters $(\hat{\mathscr{G}}_k, \hat{\theta}_k)$ are known, then the membership probability functions defined in Equation 1 and Equation 2 are identical:

$$\begin{aligned}\phi(X_V, X_C \mid k) &= \frac{\gamma_k \cdot P\left(X_V, X_C \mid \hat{\mathscr{G}}_k, \hat{\theta}_k\right)}{\sum_{k'=1}^{N} \gamma_{k'} \cdot P\left(X_V, X_C \mid \hat{\mathscr{G}}_{k'}, \hat{\theta}_{k'}\right)} \\ &= \frac{\gamma_k \cdot P\left(X_C \mid \hat{\mathscr{G}}_k, \hat{\theta}_k\right) \cdot P\left(X_V \mid X_C, \hat{\mathscr{G}}_k, \hat{\theta}_k\right)}{\sum_{k'=1}^{N} \gamma_{k'} \cdot P\left(X_C \mid \hat{\mathscr{G}}_{k'}, \hat{\theta}_{k'}\right) \cdot P\left(X_V \mid X_C, \hat{\mathscr{G}}_{k'}, \hat{\theta}_{k'}\right)} \\ &= \frac{\gamma_k \cdot P\left(X_V \mid X_C, \hat{\mathscr{G}}_k, \hat{\theta}_k\right)}{\sum_{k'=1}^{N} \gamma_{k'} \cdot P\left(X_V \mid X_C, \hat{\mathscr{G}}_{k'}, \hat{\theta}_{k'}\right)} = \tilde{\phi}(X_V \mid k)\end{aligned}$$

Following Blackwell-Rao theorem, this implies that the variance of Equation 1 is lower than the variance in Equation 2, since the estimation of $P(X_C \mid \hat{\mathscr{G}}_k, \hat{\theta}_k)$ introduces further variance in the estimation. In addition, applying our causal knowledge about the direction of the causal pathways allows us to learn the DAGs $\hat{\mathscr{G}}_k$ more accurately, reducing the variance in $P(X_V \mid \hat{\mathscr{G}}_k, \hat{\theta}_k)$. If the covariates are excluded from the analysis (Equation 3), they act as hidden variables in the probability distribution of the mutations, leading to a lower accuracy in the cluster assignment.

We will now discuss two alternative scenarios to the assumption of a constant probability distribution of the covariates across the clusters. First, instead of being constant, the probability distribution of the covariates could depend on the clusters, i.e. $\exists k, k' \in \{1, ..., K\} : P(X_C \mid k) \neq P(X_C \mid k')$. In this case, the covariates would be cluster-dependent covariates that are informative for the cluster assignment. Thus, the covariates should be treated like cluster-dependent covariates without adjustments. Another scenario could be that the probability distribution of the covariates depends on a different grouping than the clusters $g \in \{1, ..., G\}$, i.e. $P(X_C \mid k, g) \neq P(X_C \mid k, g')$. In this case, adjusting for the covariates helps to remove the bias of the other grouping in the clustering of the mutations.

**Algorithm 1:** Covariate-Adjusted Clustering

**Input:** A matrix of variables $X_V$ and a matrix of covariates $X_C$, a cutoff limit $\varepsilon$
**Output:** Cluster membership probabilities $\tilde{\phi}(X_V)$ and respective Bayesian networks $(\mathscr{G}_k, \theta_k)$

Initialize membership probabilities $\tilde{\phi}(X_V \mid k)$
**repeat**
  $i \leftarrow 0$
  Create copy of membership probabilities $\tilde{\phi}_{\text{previous}}(X_V \mid k) \leftarrow \tilde{\phi}(X_V \mid k)$
  Learn the DAGs $\mathscr{G}_k$ given the membership probabilities $\tilde{\phi}(X_V \mid k)$
  **repeat**
    $i \leftarrow i+1$
    M-Step: Learn the parameters $\theta_k$ given $\mathscr{G}_k$ and $\tilde{\phi}(X_V \mid k)$
    E-Step: Update membership probability: $\tilde{\phi}(X_V \mid k) \leftarrow \frac{\gamma_k \cdot P\left(X_V|X_C, \hat{\mathscr{G}}_k, \hat{\theta}_k\right)}{\sum_{k'=1}^{N} \gamma_{k'} \cdot P\left(X_V|X_C, \hat{\mathscr{G}}_{k'}, \hat{\theta}_{k'}\right)}$
    Update the cluster weights $\gamma_k$: $\gamma_k \leftarrow \frac{\sum_{1=1}^{N} \tilde{\phi}(X_V|k)}{N}$
  **until** $i = 10$;
  Quantify change in membership probabilities $\delta \leftarrow \sum_{X_V} \left(\tilde{\phi}(X_V \mid k) - \tilde{\phi}_{\text{previous}}(X_V \mid k)\right)^2$
**until** $\delta < \varepsilon$;

#### C.2 Algorithm

As outlined in Algorithm 1, we clustered the mutational profiles and clinical covariates using a mixture model. We used a Bernoulli mixture model to initialize the membership probabilities and learned the DAGs $\mathscr{G}_k$ with corresponding local probability distributions $\theta_k$ using the **BiDAG** package (GPL-3)[3]. Due to the high computational cost of the DAG structure search, we only relearned the DAGs $\mathscr{G}_k$ for every tenth update of the membership probabilities.

Our computations were performed on one CPU core of the AMD EPYC 7H12 processor (2.6 GHz nominal, 3.3 GHz peak) and 256 GB of DDR4 memory clocked at 3200 MHz. Code implementing our method is publicly available as a software package at `https://CRAN.R-project.org/package=clustNet`.

While our algorithm presents several advantages, it comes with the trade-off of higher computational cost compared to traditional clustering methods like k-means or Bernoulli mixture models. The computational cost scales notably with an increase in the number of variables. In our simulations, the computation time averaged 4.18 minutes over 20 repetitions with the following parameters: 4 clusters, 20 variables, 5 covariates, and 4800 samples. Clustering the mutational and clinical profiles from the pan-myeloid data (9 clusters, 59 variables, 13 covariates, 1323 samples), our method took 3.4 hours to converge. The clustering of the TCGA data (22 clusters, 201 variables, 24 covariates, 8085 samples) took 30.32 hours to converge. By parallelizing the structure learning step of the individual clusters, we were able to reduce the runtime of our algorithm by a factor of 1.6 for two clusters.

#### C.3 Evolutionary Perspective

Cancer progression can be described as the somatic evolution of mutations in the genome[?]. The occurrence of mutations is commonly modelled using mutation trees, in which the presence of one mutation makes another mutation more likely to occur[?]. This process of genetic progression creates mutational patterns of probabilistic relationships among the genes. Bayesian networks are a natural framework to model the mutational patterns that are created through somatic evolution[?]. Bayesian networks are a powerful tool to model the probabilistic relationships amongst random variables and allow for a natural integration of covariates. While an evolutionary tree describes possible orders in which the mutations can occur[?], a Bayesian network can describe more general probabilistic relationships amongst the mutations[24]. Note that the nodes in the mutation tree contain a list of mutated genes, whereas the nodes in the Bayesian network contain the genes themselves, which can either be mutated or not.

Besides Bayesian networks, there are various other probabilistic graphical models. The most popular models include tree graphical models and Markov random fields (undirected graphical models)[?]. However, mutational trees induce nodes with more than one parent, which can not be modelled by tree graphical models. On the other hand, undirected models can model evolutionary trees, but can not model the order in which the genes occur.

Clinical covariates such as sex and age are closely related to the mutations in specific genes[4,26]. Such dependencies can be efficiently modelled in the framework of Bayesian networks by edges from the covariates to the respective genes.

#### D Number of clusters

To determine the optimal number of clusters, we clustered the data using a range of cluster numbers and hyperparameter $\chi$. We used the AIC score as a criterion for selecting the best-fitting number of clusters, with lower scores indicating better fits. In this analysis, we neglected the edges between nodes, which is equivalent to using a Bernoulli mixture model. We focused on mutations and cytogenetic variables to determine the number of clusters as we were interested in cluster associations based on genetic patterns. Including the clinical covariates would lead to a higher number of clusters, as these variables are not as sparse as the mutational and cytogenetic profiles. While a higher number of clusters may also yield meaningful cluster associations, it reduces the number of samples per cluster, resulting in lower certainty in the learned networks. Applying our method to a larger dataset may therefore allow learning a higher number of clusters.

Figure S4 shows the AIC scores for different numbers of clusters and hyperparameters $\chi$, where the difference to the highest value is displayed; cluster assignments that contained empty clusters are marked in grey. We chose the cluster number nine, as it most frequently resulted in the highest AIC score, which varied across the range of $\chi$. For larger values of $\chi > 2$, the individual data points had a lower weight, leading to a smaller number of clusters.

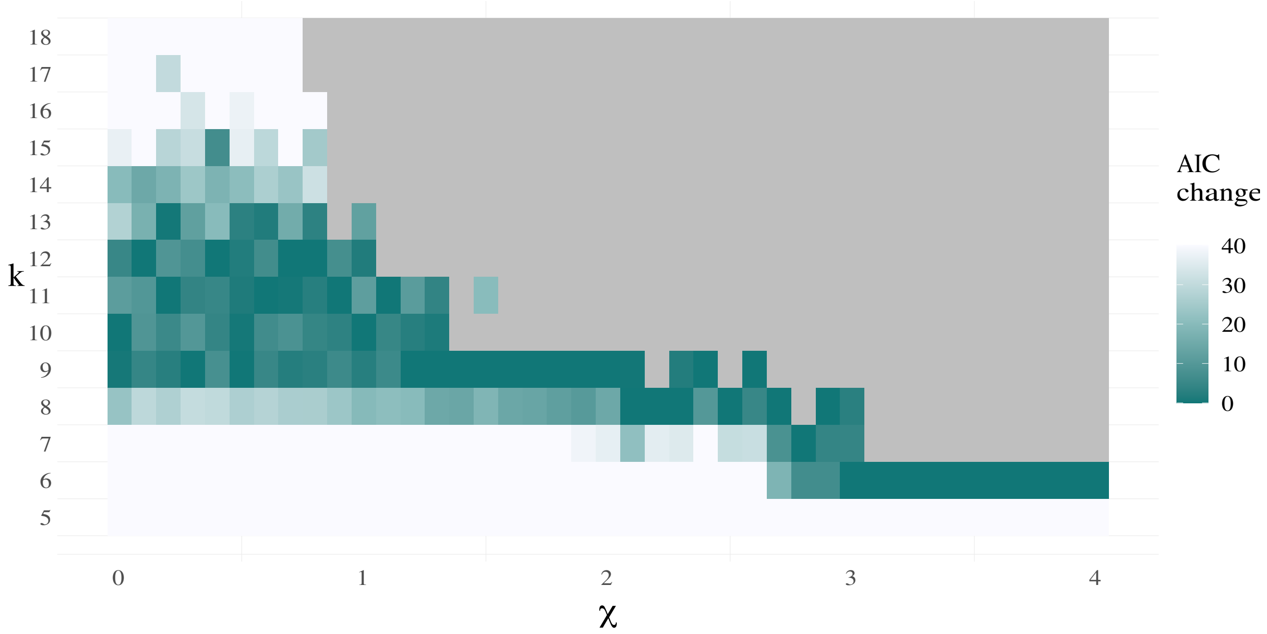


**Figure S4.** AIC change over different numbers of clusters. Grey areas mark the configurations where empty clusters were present.

**Table S2.** Summary table of patient characteristics.

| Characteristic | All | AML | MDS | CMML | MPN |
|---|---|---|---|---|---|
| Number of patients | 1323 | 543 | 492 | 118 | 170 |
| Median age (years) | 66 | 62 | 69 | 71 | 59 |
| Range of age (years) | 18-94 | 18-94 | 20-91 | 51-92 | 18-85 |
| Number of males | 792 | 296 | 324 | 87 | 85 |
| Number of females | 531 | 247 | 168 | 31 | 85 |

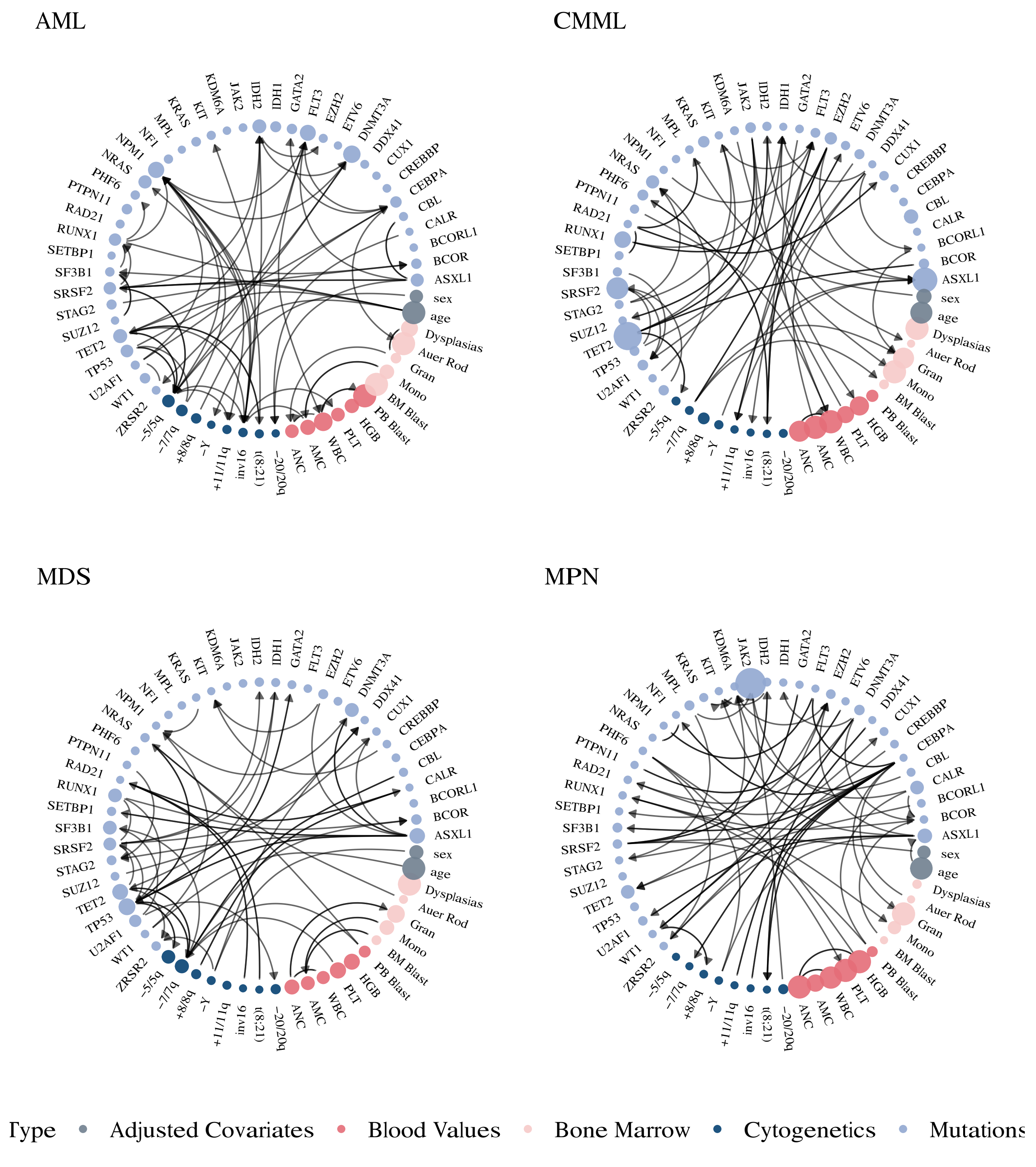


**Figure S5.** Cancer-specific networks of the clusters A-I, reproducing Figure 2 for the full set of variables. Genomic features are illustrated in blue, blood and bone marrow values in red and demographic variables in grey. The node size for mutations and cytogenetic variables corresponds to their frequency within each cluster, with more frequently mutated genes appearing as larger nodes. Conversely, the node size of continuous variables gets larger when the values of those variables are higher.

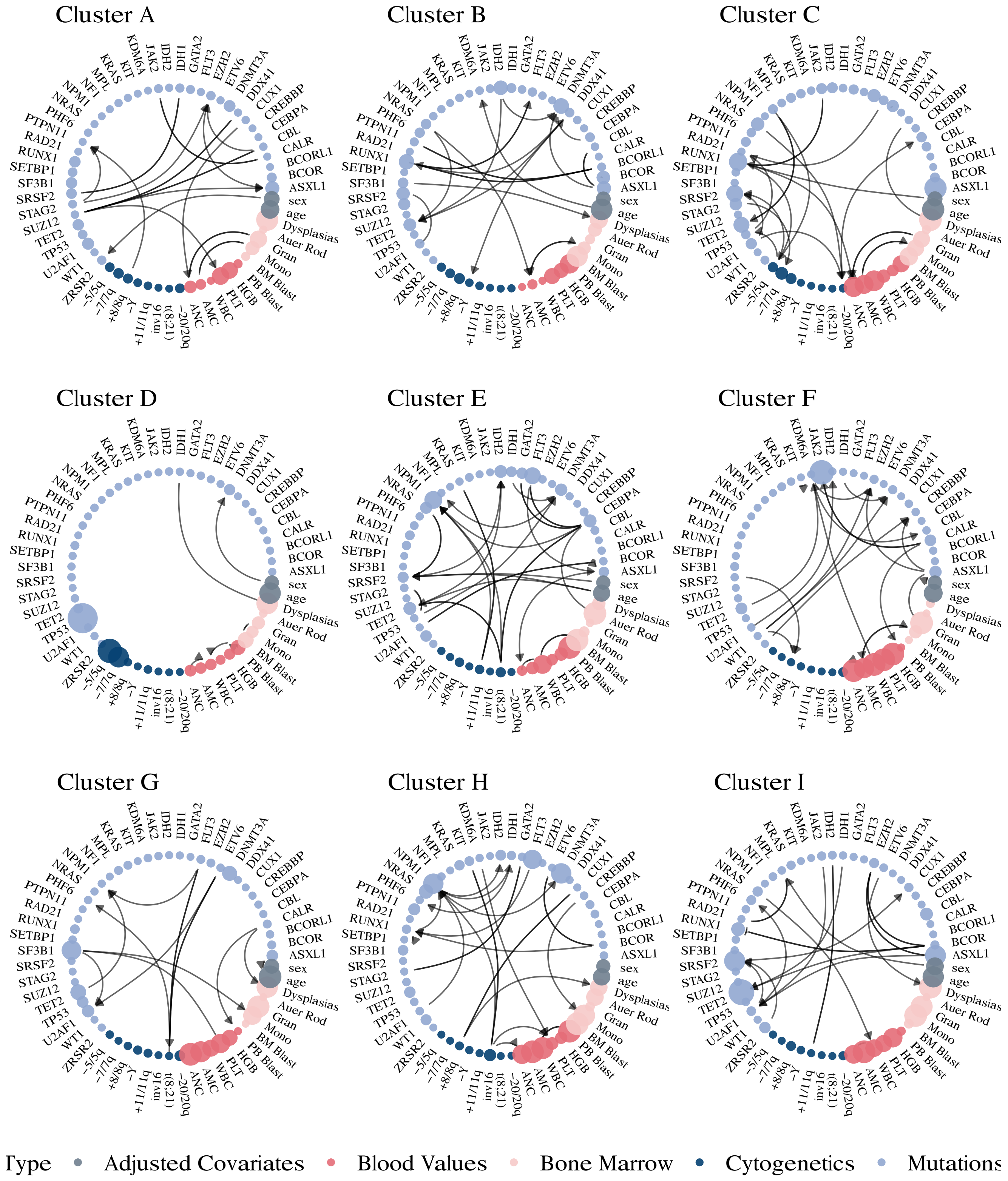


**Figure S6.** Cluster-specific networks of the clusters A-I, reproducing Figure 4 for the full set of variables. Genomic features are illustrated in blue, blood and bone marrow values in red and demographic variables in grey. The node size for mutations and cytogenetic variables corresponds to their frequency within each cluster, with more frequently mutated genes appearing as larger nodes. Conversely, the node size of continuous variables gets larger when the values of those variables are higher.

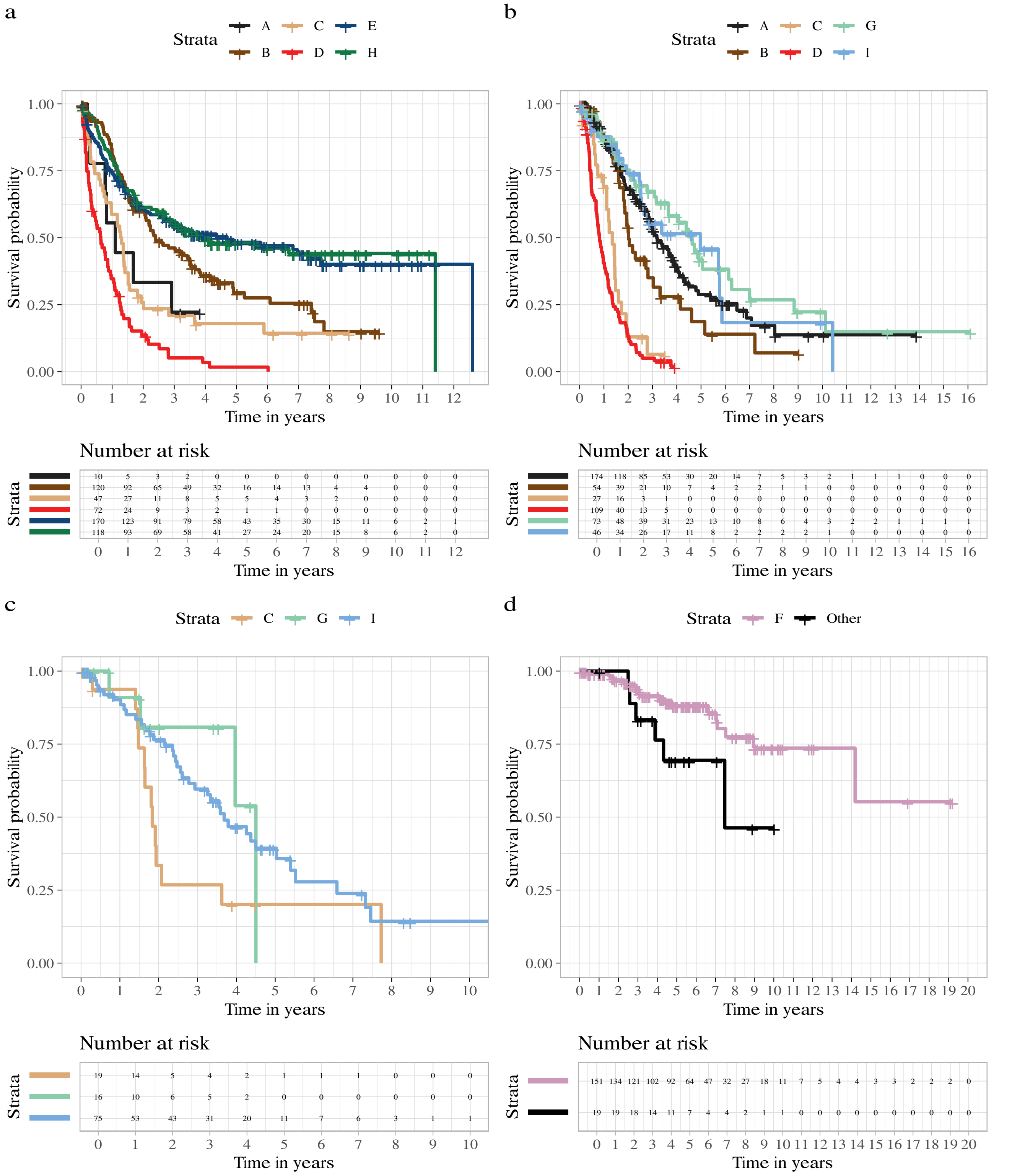


**Figure S7.** Kaplan-Meier survival curves for AML (a), MDS (b), CMML (c), and MPN (d) patients, stratified by cluster. Groups with fewer than 10 individuals are omitted for clarity. In MPN, due to only one cluster with more than 10 individuals, other clusters are grouped together.

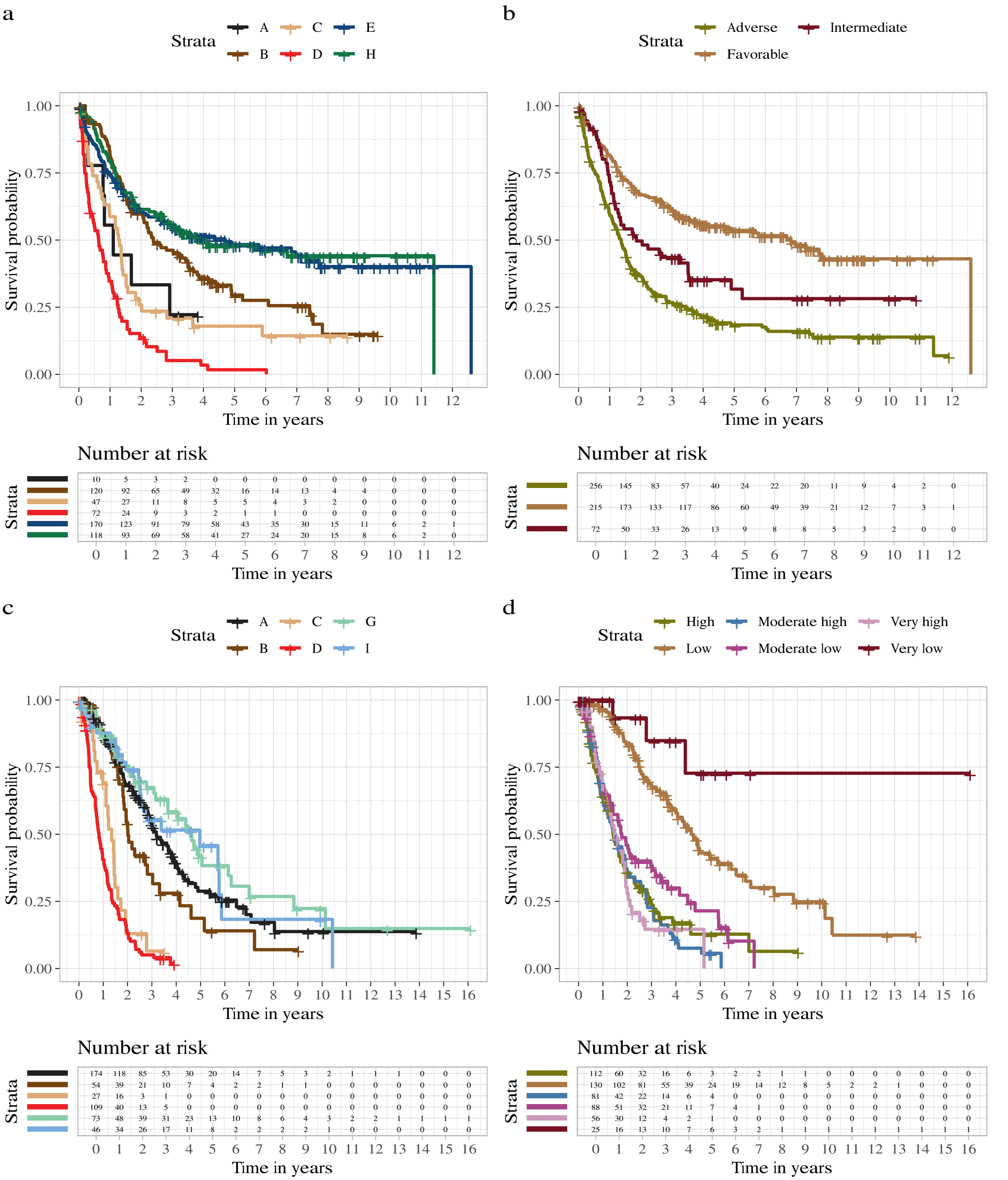


**Figure S8.** Kaplan-Meier survival curves comparing distinct clusters and ELN2022 and IPSS-M risk classifications in AML and MDS, respectively. Displayed are the survival probabilities for distinct clusters in AML patients (a) and their survival outcomes stratified by ELN2022 risk classification (b). Corresponding analyses for MDS include survival probabilities across distinct clusters (c) and according to IPSS-M risk classification (d). Groups with fewer than 10 individuals are omitted for clarity.

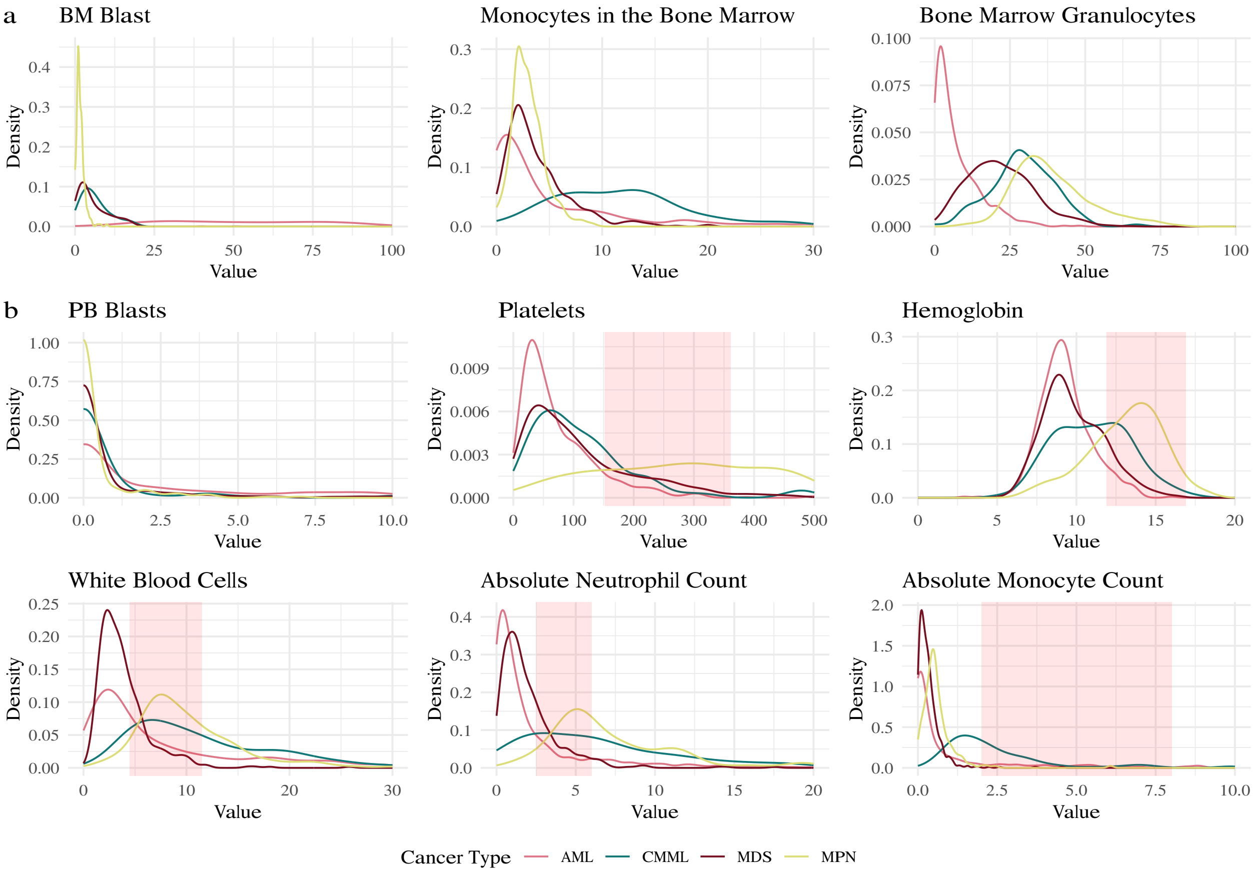


**Figure S9.** Distribution of the continuous variables in bone marrow (a) and blood (b). Displayed are the kernel density plots of the distributions for each cancer type. The normal range observed in healthy individuals is displayed in transparent red.

**Table S3.** Summary of variables contained in our patient cohort. Excluded variables contain genes with a prevalence below 1% and clinical information that does not display differences across the disease types.

| Variable type | Variable name |
|---|---|
| Demographic Variables | Age, Gender |
| Clinical Information | Cancer type, ANC, AMC, WBC, PLT, HGB, PB Blast, BM Blast, Mono, Gran, Auer Rod, Dysplasias, ALC |
| Mutations | *ASXL1, BCOR, BCORL1, BRAF, CALR, CBL, CEBPA, CREBBP, CSF3R, CUX1, DDX41, DNMT3A, EED, ETNK1, ETV6, EZH2, FBXW7, FLT3, GATA2, GNAS, IDH1, IDH2, IKZF1, IL7R, JAK1, JAK2, JAK3, KDM6A, KIT, KRAS, MAP2K1, MLL, MPL, NF1, NOTCH1, NPM1, NRAS, PAX5, PHF6, PIGA, PTPN11, RAD21, RUNX1, SETBP1, SF3B1, SH2B3, SMC1A, SMC3, SRSF2, STAG2, STAT3, SUZ12, TET2, TP53, U2AF1, U2AF2, WT1, ZRSR2* |
| Cytogenetics | -5/5q, -7/7q, +8/8q, -Y, +11/11q, inv16, t(8;21), -20/20q |
| Excluded variables | *BRAF, CSF3R, EED, ETNK1, FBXW7, GNAS, IKZF1, IL7R, JAK1, JAK3, MAP2K1, MLL, NOTCH1, PAX5, PIGA, SH2B3, SMC1A, SMC3, STAT3, U2AF2*, ALC |

**Table S4.** Summary table of survival analysis by each cluster and cancer type. Groups with fewer than 10 individuals are omitted for clarity.

| Cluster | Cancer type | Records | Events | RMST | SE(RMST) | Median OS | 95LCL | 95UCL |
|---|---|---|---|---|---|---|---|---|
| A | Overall | 191 | 94 | 5.4 | 0.7 | 3.2 | 2.8 | 3.9 |
| B | Overall | 175 | 113 | 4.9 | 0.6 | 2.3 | 2.0 | 3.2 |
| C | Overall | 98 | 75 | 3.6 | 0.7 | 1.4 | 1.3 | 1.6 |
| D | Overall | 182 | 165 | 1.0 | 0.1 | 0.8 | 0.6 | 0.9 |
| E | Overall | 173 | 88 | 6.4 | 0.5 | 4.7 | 2.9 | |
| F | Overall | 158 | 21 | 14.4 | 1.1 | | 14.2 | |
| G | Overall | 98 | 42 | 6.7 | 1.0 | 4.6 | 4.0 | 7.0 |
| H | Overall | 121 | 62 | 6.1 | 0.5 | 3.9 | 2.7 | |
| I | Overall | 127 | 65 | 5.1 | 0.7 | 3.6 | 2.8 | 5.4 |
| A | AML | 10 | 7 | 5.1 | 2.5 | 1.1 | 0.8 | |
| A | MDS | 174 | 85 | 5.4 | 0.7 | 3.2 | 2.8 | 4.0 |
| B | AML | 120 | 79 | 5.3 | 0.8 | 2.4 | 2.1 | 3.5 |
| B | MDS | 54 | 33 | 3.8 | 0.9 | 2.0 | 1.8 | 3.3 |
| C | AML | 47 | 38 | 3.8 | 1.0 | 1.3 | 1.0 | 1.5 |
| C | CMML | 19 | 13 | 3.0 | 0.6 | 1.8 | 1.6 | |
| C | MDS | 27 | 22 | 2.4 | 1.0 | 1.3 | 1.1 | 1.7 |
| D | AML | 72 | 68 | 1.0 | 0.1 | 0.6 | 0.4 | 1.0 |
| D | MDS | 109 | 96 | 1.4 | 0.3 | 0.8 | 0.7 | 1.0 |
| E | AML | 170 | 87 | 6.4 | 0.5 | 4.7 | 2.7 | |
| F | MPN | 151 | 19 | 14.7 | 1.1 | | 14.2 | |
| G | CMML | 16 | 4 | 3.7 | 0.4 | 4.5 | 4.0 | |
| G | MDS | 73 | 34 | 6.4 | 1.1 | 4.6 | 3.7 | 7.0 |
| G | MPN | 8 | 3 | 10.7 | 3.5 | 7.5 | 7.5 | |
| H | AML | 118 | 61 | 6.1 | 0.5 | 3.9 | 2.7 | |
| I | CMML | 75 | 39 | 4.5 | 0.5 | 3.7 | 2.9 | 5.5 |
| I | MDS | 46 | 22 | 4.7 | 0.7 | 5.0 | 2.6 | |

**Table S5.** Summary table of survival analysis by each risk classification and cancer type.

| Risk | Risk score | Records | Events | RMST | SE(RMST) | Median OS | 95LCL | 95UCL |
|---|---|---|---|---|---|---|---|---|
| Adverse | ELN2022 (AML) | 256 | 198 | 3.3 | 0.4 | 1.4 | 1.1 | 1.5 |
| Favorable | ELN2022 (AML) | 215 | 102 | 6.9 | 0.4 | 6.8 | 3.9 | |
| Intermediate | ELN2022 (AML) | 72 | 45 | 5.8 | 0.9 | 1.9 | 1.3 | 3.5 |
| Very High | IPSS-M (MDS) | 56 | 39 | 1.9 | 0.2 | 1.5 | 1.1 | 1.9 |
| High | IPSS-M (MDS) | 112 | 77 | 2.8 | 0.6 | 1.4 | 1.3 | 1.8 |
| Moderate High | IPSS-M (MDS) | 81 | 62 | 2.0 | 0.2 | 1.5 | 1.1 | 2.0 |
| Moderate Low | IPSS-M (MDS) | 88 | 56 | 2.8 | 0.3 | 1.8 | 1.4 | 3.0 |
| Low | IPSS-M (MDS) | 130 | 57 | 6.2 | 0.6 | 4.6 | 3.8 | 6.5 |
| Very Low | IPSS-M (MDS) | 25 | 3 | 12.6 | 1.8 | | 4.4 | |

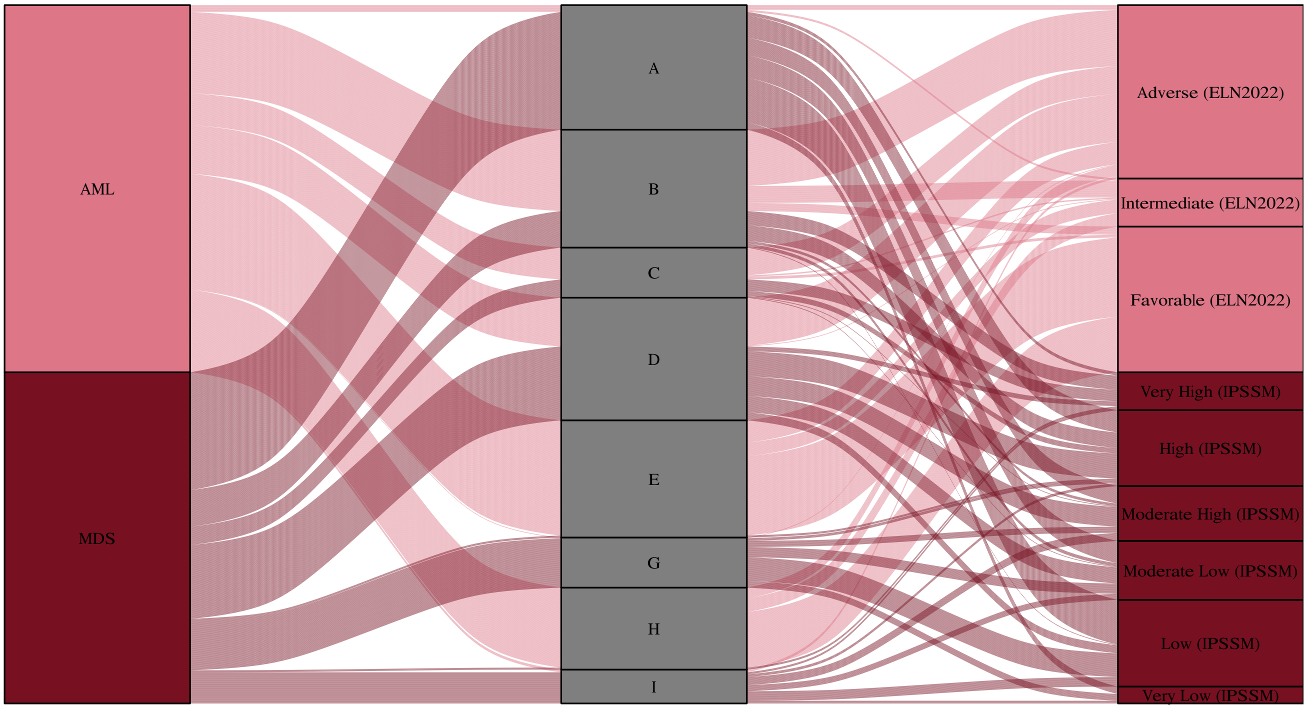


**Figure S10.** Sankey plot of AML and MDS risk scores. Clusters G and I are not displayed as they primarily contain CMML and MPN samples.

**Table S6.** Likelihood ratio for our novel subgroups compared to those generated by the ELN2022 and IPSS-M risk scores. The table includes corrected likelihood ratios, which account for age, sex, and cancer type in the survival analysis. The final two rows offer further corrections, incorporating the effects of the aforementioned risk scores into the survival analysis. In our cluster analysis and risk score calculations, we did not include certain features due to their absence in our dataset. Specifically, the following were excluded: complex karyotype, -17/del(17p), *TP53*$^{\text{multi-hit}}$, *GNB1, PPM1D, PRPF8*, t(9;11), t(6;9), t(9;22), inv(3), and i(17).

| Grouping | Cancer type | Corrected likelihood ratio | Likelihood ratio | P-value |
|---|---|---|---|---|
| CANclust | AML | **29.2** | **165.1** | $9.0 \times 10^{-10}$ |
| ELN2022 | AML | 16.9 | 140.6 | $4.2 \times 10^{-5}$ |
| CANclust | MPN | **76.4** | **154.4** | $5.2 \times 10^{-29}$ |
| IPSS-M | MPN | 53.9 | 114.1 | $1.1 \times 10^{-19}$ |
| CANclust (beyond ELN2022) | AML | 21.2 | 182.9 | $1.2 \times 10^{-6}$ |
| CANclust (beyond IPSS-M) | MDS | 53.9 | 221.8 | $1.1 \times 10^{-19}$ |

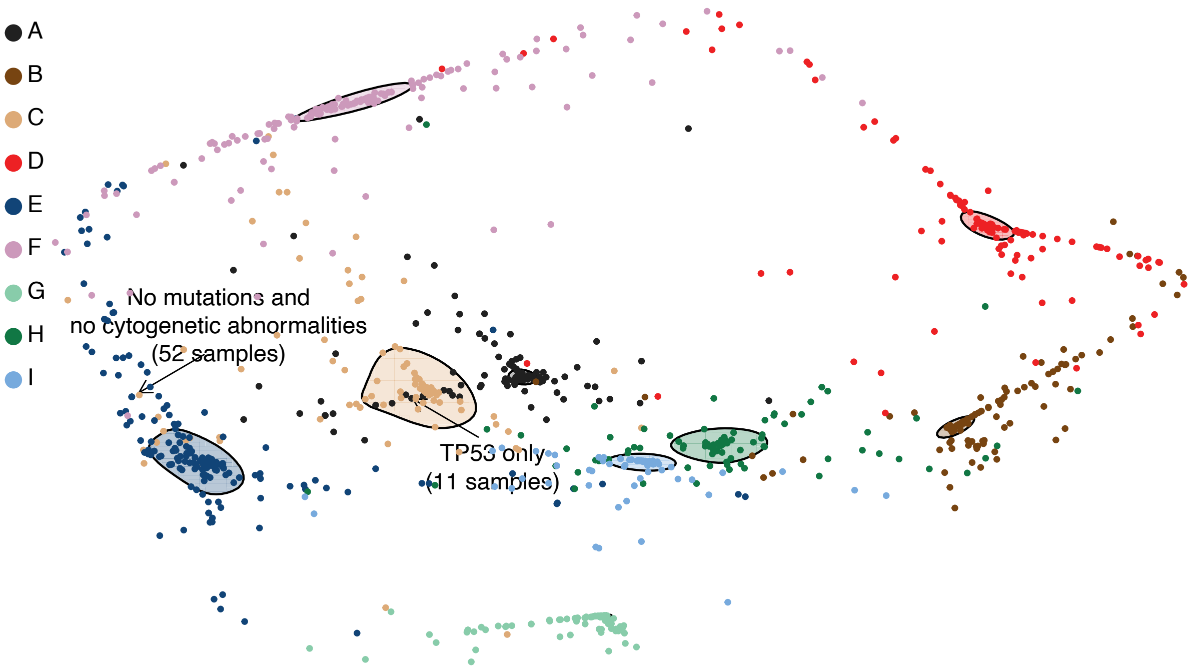


**(a)** Density plot of the different clusters coloured by cluster association.

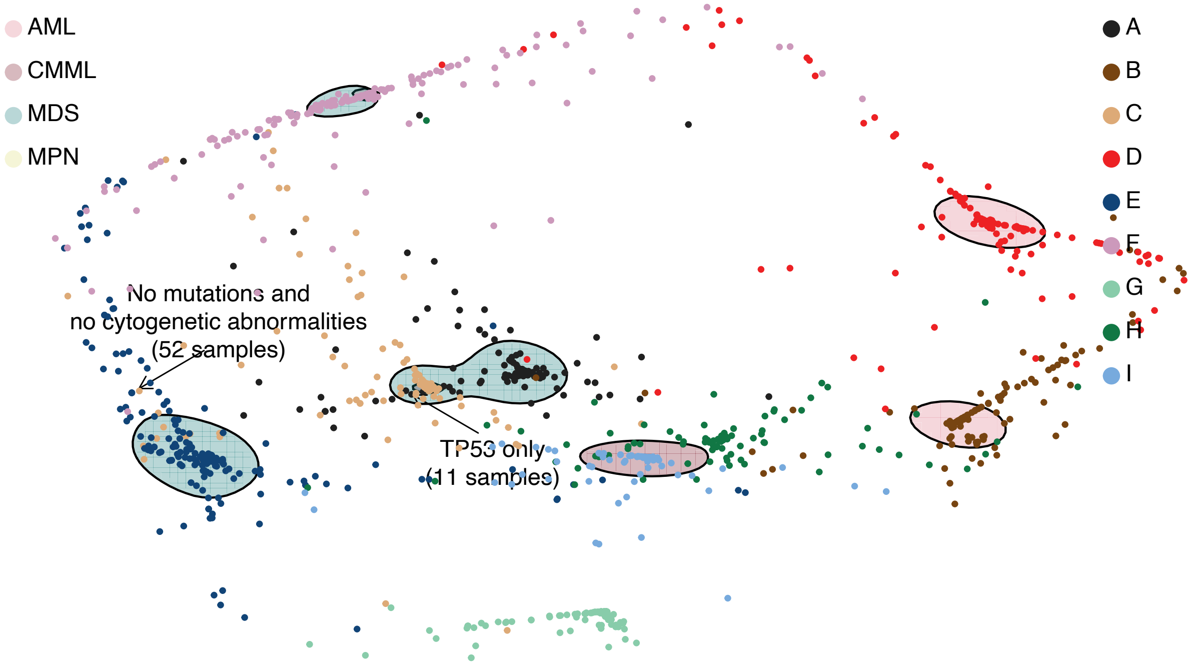


**(b)** Density plot of the different cancer types coloured by cancer type and cluster.

**Figure S11.** Visualized in 2D are the patient samples based on how well they fit to each cluster-specific Bayesian network. Each dot represents a patient and is coloured by its corresponding cluster. Solid shapes depict contours that collectively encompass 50% of the corresponding cluster (a) or cancer type (b).

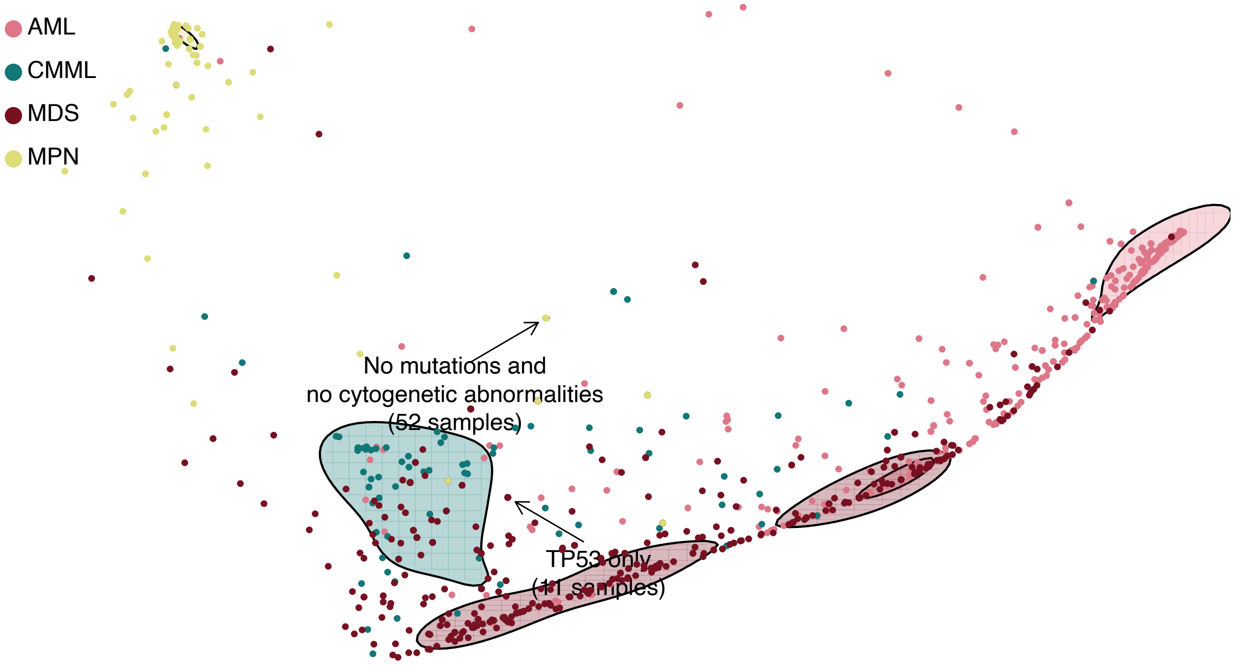


**(a)** Density plot of the different cancer types coloured by cancer type.

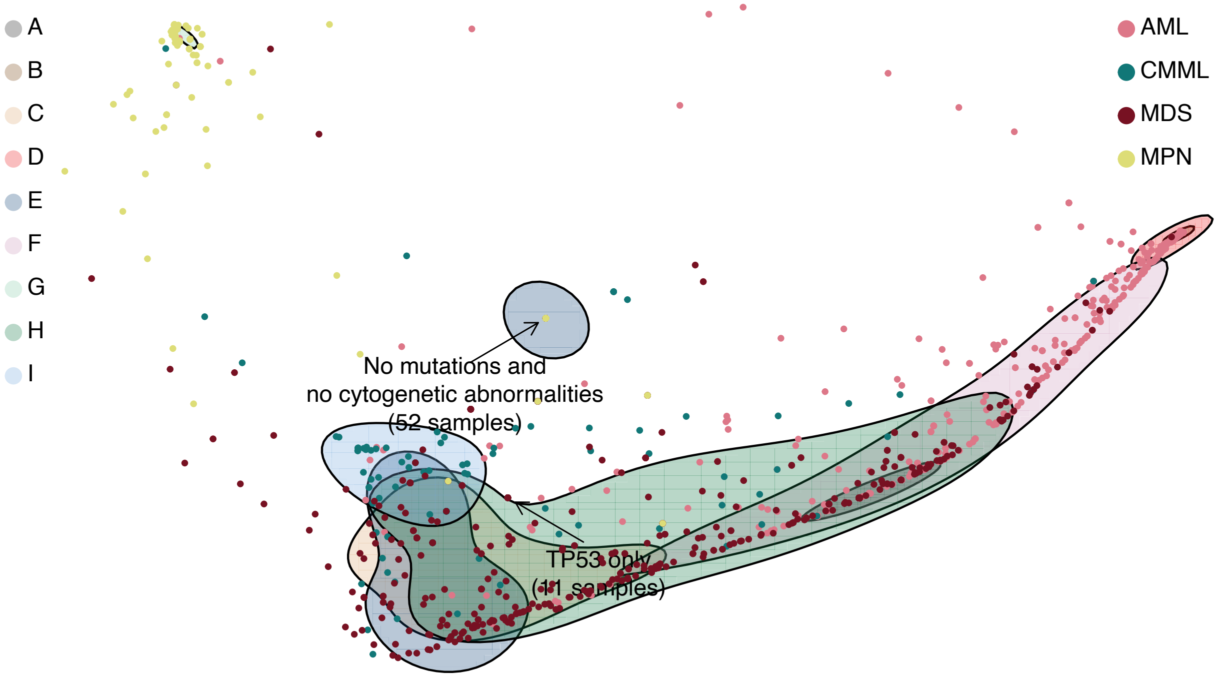


**(b)** Density plot of the different cancer types coloured by cluster and cancer type.

**Figure S12.** Visualized in 2D are the patient samples based on how well they fit to each cancer type specific Bayesian network. Each dot represents a patient and is coloured by its corresponding cancer type. Solid shapes depict contours that collectively encompass 50% of the corresponding cancer type (a) or cluster (b).
